# Anatomically Informed 3D Printed CT phantoms: The First Step of a Pipeline To Identify Robust Quantitative Radiomic Features

**DOI:** 10.1101/773879

**Authors:** Usman Mahmood, Aditya Apte, Christopher Kanan, David D.B. Bates, Giuseppe Corrias, Lorenzo Manneli, Jung Hun Oh, Yusuf E. Erdi, John Nguyen, Joseph O. Deasy, Amita Shukla-Dave

## Abstract

**Purpose:** This study investigates the robustness of quantitative radiomic features derived from computed tomography (CT) images of a novel patient informed 3-D printed phantom, which captures the morphological heterogeneity of tumors and normal tissue observed on CT scans.

**Methods:** Using a novel voxel-based multi-material three-dimensional (3D) printer, an anthropomorphic phantom that was modeled after diseased tissue seen on 6 patient CT scans was manufactured. Four patients presented with pancreatic adenocarcinoma tumors (PDAC), 1 with non-small cell lung carcinoma (NSCLC) and 1 with advanced stage hepatic cirrhosis. The 5 tumors were segmented, extracted and then imbedded into CT images of the heterogenous portion of the cirrhotic liver. The composite scan of the implanted tumor within the background cirrhotic liver was then 3D printed. The resultant phantom was scanned sequentially, 30 times with a clinical CT scanner using a reference CT protocol. One hundred and four quantitative radiomic features were then extracted from images of each lesion to determine their repeatability. Repeatability of each radiomic feature was evaluated using the within subject coefficient of variation (wCV, %). A feature with a wCV (%) > 10% was considered as being unrepeatable. A subset of the repeatable features that were also found to be prognostic for lung and pancreatic cancers were then assessed for their percent deviation (pDV, %) from reference values. The reference values were those derived from the repeatability portion of this study. The assessment was conducted by re-scanning the phantom with 11 different clinically relevant sets of scanning parameters. Deviation of radiomic features derived from images of each tumor across all sets of scanning parameters was assessed using the percent deviation relative to the reference values.

**Results:** Twenty nine of the 104 features presented with wCV (%) > 10%. The lack of repeatability was found to depend on tumor type. The only class of radiomic features with a wCV (%) < 10% were those calculated using the neighboring grey level dependence-based matrices (NGLDM). Notably, skewness, first information correlation, cluster shade, Haralick correlation, autocorrelation, busyness, complexity, high gray level zone emphasis, small area high gray level emphasis, large area low gray level emphasis, large area high gray level emphasis, short run high grey level emphasis, and valley radiomic features had wCV (%) values > 10% for select tumors within the phantom. Two radiomic features prognostic for NSCLC, energy and grey level non-uniformity, had pDV’s (%) that exceeded 30% across all scanning techniques. The pDV (%) for the 4 radiomic features prognostic for PDAC tumors depended on tumor type and selected scanning parameter. Application of the lung kernel caused the largest pDV’s (%). Scans acquired with the reduced tube current of 100 mA and reconstructed with the bone kernel yielded pDV’s (%) within ± 10%.

**Conclusion:** We demonstrated the feasibility with which patient informed 3D printed phantoms can be manufactured directly from lesions seen on CT scans, and demonstrate their potential use for the assessment of robust quantitative radiomic features.

## Introduction

Quantitative radiomic features have emerged as an objective means to characterize the degree of inter and intratumor heterogeneity seen on medical imaging exams, such as those from computed tomography (CT) scans [1]. There are several ways in which radiomic analysis may be useful in diagnostic medical imagine. For example, establishing a link between radiomic features and gene expression patterns may provide imaging biomarkers that can be incorporated into precision medicine models for patients who undergo imaging with CT [2, 3]. The ability to quantitatively analyze tumors with the same histological subtype using CT scans could enhance diagnostic imaging, potentially assisting with the development of targeted therapies[4-6].

However, the sensitivity of quantitative radiomic features depend on the attributes of each CT system [7-9]. Previous attempts to characterize the uncertainty resulting from different CT acquisition parameters have largely involved uniform, homogenously textured or unrealistically shaped tumors embedded into phantoms. These unrealistic representations of human tissue may over or understate the uncertainty of quantitative radiomic features extracted from tumors seen on CT scans[10]. For example, the heterogeneity of tissue surrounding a lesion is known to influence the local noise and resolution properties of the identified structures observed on CT scans, especially when iterative reconstruction algorithms are used [11-13]. Uniform phantoms are incapable of capturing the degree to which the unique attributes of each CT scanner influence quantitative radiomic features [7, 10, 14-16]. Hence, the acceptable range of variation and the reporting of repeatability and reproducibility may be falsely stated [8, 17-19].

In contrast to the approaches summarized above, we develop a novel three-dimensional (3D) printed radiomic phantom that replicates the morphology and reproduces the contrast differences of diseased human tissue seen on CT exams. With the use of a multi-material 3D printer, we propose a method to generate realistic patient informed phantoms that can be used to quantify the robustness of radiomic features derived from patient CT scans.

Our specific aims are to use a novel voxel-based multi-material 3-D printing technology to i) design, fabricate and develop the first iteration of an anatomically relevant 3-D printed phantom for radiomic robustness analysis, and ii) test the radiomic features derived thereafter for repeatability and reproducibility relative to reference values.

## Methods

The specific aspects of this study included the manufacturing process of the patient informed 3D printed phantom. Second, the similarity between the 3D printed phantom with the human tissue seen in the DICOM CT scans it was modeled after was assessed. Third, quantitative radiomic features derived from the 3D printed phantom were tested for repeatability. Lastly, the deviation of a subset of radiomic features found to be repeatable and prognostic were assessed relative to reference values from the repeatability part of the study.

### 1.1 3D Printed Phantom Fabrication

The multi-material 3D printer and associated software, Voxel Print, (PolyJet Objet 260 Connex 3, Stratasys, Eden Prairie, Minnesota) allows for the deposition of droplets of ultraviolet-curable photopolymer resins in a layer by layer inkjet like printing process [14]. The droplet resolution is 600 and 300 dots per inch (DPI), with a slice thickness of 30 *μ*m. This exceeds the resolution of a typical C^i^T scanner (0.625 × 0.625 × 0.625 mm) [14]. As a result, the morphological detail of tumors seen on CT scans may be preserved, while the heterogenous contrast differences of tumors may be reproduced with some degree of precision[11, 20, 21]. Similar to Bader et al. [20], a graphical overview of the methods used are shown in Figure 1. Displayed in Figure 1a is a cross-sectional CT scan of a single patient, with non-small cell lung carcinoma (NSCLC), from the publicly available test-retest CT scans of the Reference Image Database to Evaluate Therapy Response (RIDER) collection hosted by the Cancer Imaging Archive (TCIA) dataset [22-24]. The lung volume (Figure 1b) with the associated tumor (Figure 1c) were used to design the first 3D print (Figure 1f), whose purpose was to physically visualize the potential of voxel-based 3D printing to be mapped and converted to resin material gradients.

**Fig. 1.**
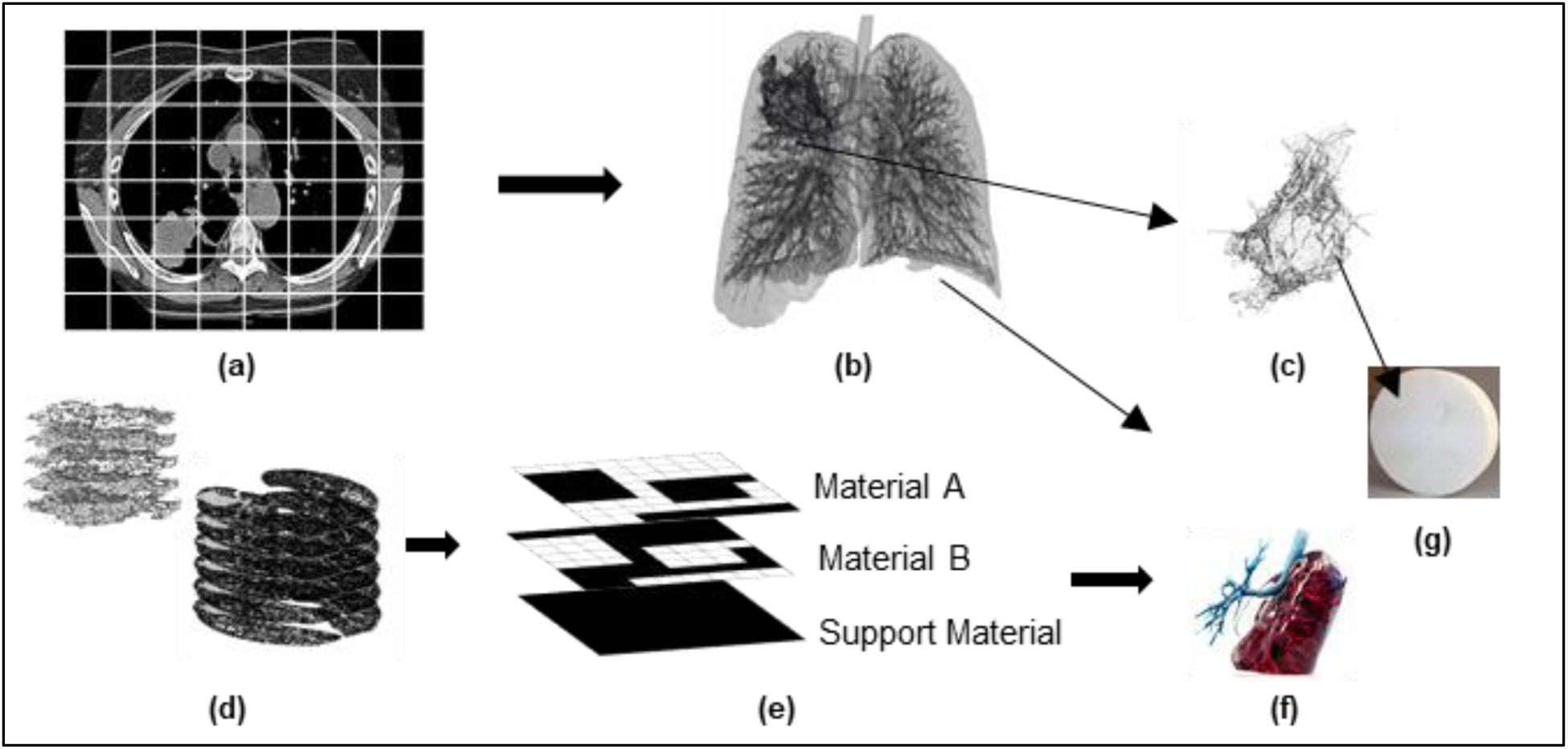
Generation workflow of an informed radiologic phantom. (a) A cross sectional slice from a patient CT scan shows a non-small cell lung carcinoma (NSCLC) lesion within the right upper lobe. (b) The volumetric representation of the segmented lung and (c) extracted tumor. The dimensions of these volumes defined the overall size of the final print. (d) The lung and tumor volumes were resliced into the resolution of the 3D printer and stacked into layers as shown. (e) Each layer from (d) was then dithered using the Floyd Steinberg dithering algorithm into binary raster files. Three sets of raster files were needed, one for each resin material. These files define the spatial location of each resin material. (f) The resulting 3D print of the lung volume. This print was designed to be a physical visualization of the entire patient lung with tumor. (g) The actual phantom used in this study. The arrow is pointing to the location of the NSCLC lung tumor.

The second 3D print included 4 pancreatic adenocarcinoma (PDAC) tumors. These tumors were manually segmented by experienced radiologists. These patients received contrast enhanced abdominal CT scans on a single 64 slice CT scanner (HD750, General Electric, Madison Wisconsin). The 5^th^ tumor was the NSCLC mass seen in Figure 1c. All segmented tumors were embedded into a heterogenous background that was modeled after a 6^th^ patient who presented with advanced stage hepatic cirrhosis on an abdominal contrast enhanced CT scan. The area and number of slices of the cirrhotic liver was dictated by the dimensions of each tumor. Placement of the tumors within the background cirrhotic liver was arbitrary.

To prepare the volumes for 3D printing, the voxel Hounsfield unit (HU) values were normalized to range from 0 to 1. The new fractional intensity values were used to dictate the proportion of resin material that would be deposited in any given voxel [11]. To obtain the gray scale intensity gradients seen in CT scans, interpolation of each volume was necessary. This was achieved by using the Whittaker–Shannon (SINC) interpolation method, where each voxel was super-sampled to the resolution of the 3D printer (Figure 1d).

Lastly, the layers were dithered using the Floyd-Steinberg dithering algorithm (Figure 1e) into binary raster files and each volume was then embedded into the cirrhotic liver background. These bitmap files defined the spatial allocation of each material to be used for 3D printing. The multi-material printer has available 3 different resin materials for printing. As a result, three sets of bitmap files were generated, one set for each resin material. Within any bitmap, a value of 1 indicates deposition of material A and a value of 0 indicates no material will be deposited. The first set of bitmaps encoded the deposition location of the resin material A (Figure 1e). Then, the material A bitmaps were inverted so that a value of 0 now had a value of 1. These inverted bitmaps encoded the allocation of resin material B (Figure 1e). The 3rd set of bitmaps consisted of all zeros since two materials with opposing densities were enough to generate the desired contrast differences. The resulting 3D print is displayed in Figure 2a. Shown in Figure 2b, c are cross sectional CT images of the proposed phantom.

**Fig. 2.**
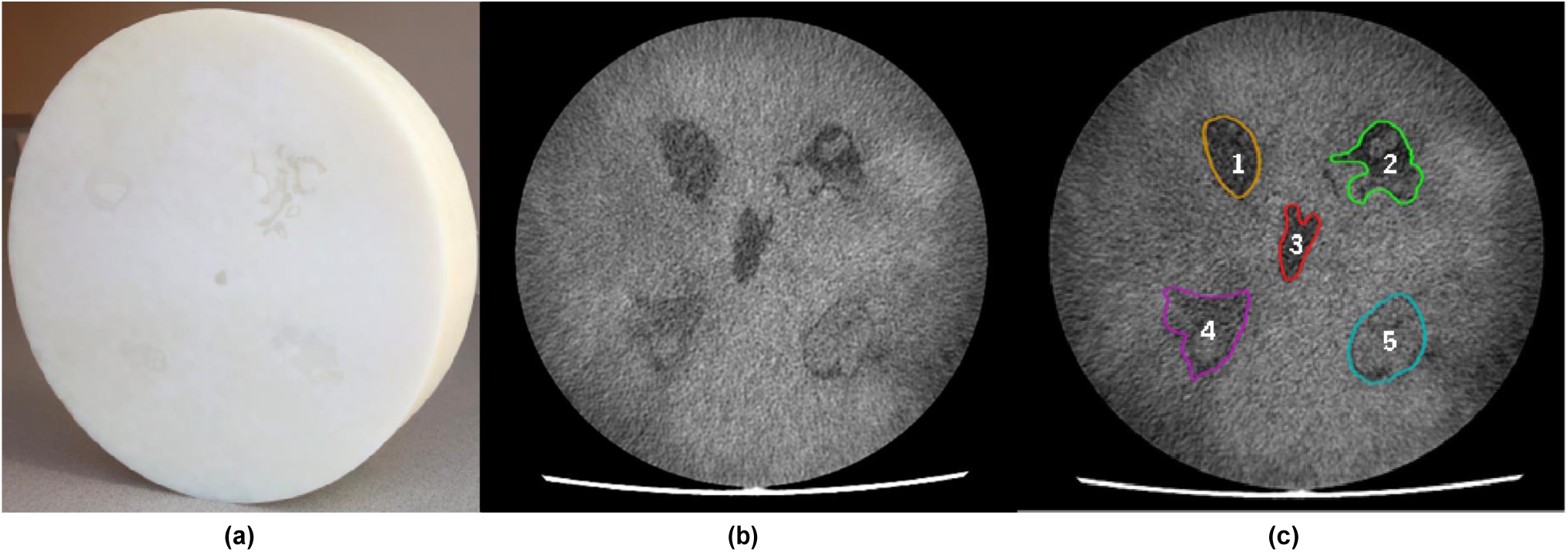
The first iteration of the 3D printed phantom used in this study. (a) The circular phantom consists of tumors embedded within a heterogenous background derived from a patient’s liver scan. (b) Axial slice generated from a computed tomography (CT) scan shows the embedded tumors within the background tissue. (c) Contours that were generated from an experienced radiologist. The tumor labels identified the tumor type: 1-non-small cell lung carcinoma (NSCLC); 2-5 are pancreatic tumors.

The two-resin material used in this study were VeroWhite (material A) and TangoPlus (material B). They were selected based on their attenuation properties observed within CT scans. The scanning parameters used were: tube potential of 120 kVp, tube current of 280 mA, filtered back projection (FBP) reconstruction algorithm with a standard kernel, 1.25 mm slice thickness with an interval of 1.25 mm. Multiple ROIs were drawn along the samples of each resin. The HU values were measured to range from 125 ± 5 HU for material A and 65 ± 5 HU for material B.

### 1.2 Repeatability and Reproducibility Scanning Parameters

A 64 slice CT scanner (HD750, General Electric, Madison Wisconsin) scanner was used to acquire 30 repeat scans of the 3D printed radiomic phantom (Figure 2a). All scans were completed without repositioning of the phantom. The scanning parameters were: 120 kVp, 280 mA, 0.7 second, pitch of 0.984, filtered back projection algorithm with a standard kernel, total collimation of 40 mm, display field of view (DFOV) 25 cm, reconstructed slice thickness and interval of 5 mm. According to Nerwell, J.D. et al. [25], during the development of quantitative CT metrics of lung disease, multiple CT vendors agreed to provide neutral reconstruction kernels that could provide comparable CT attenuation values. For the CT vendor used in this study, the neutral scanning protocols consisted of those that used filtered back projection algorithms with the standard kernel. Consequently, radiomic features extracted from this protocol were considered as reference values.

Since CT scanners come equipped with several user adjustable scan options, additional images were acquired using 7 different reconstruction algorithms, three different kernels, a larger voxel size, a reduced tube potential (kVp) and reduced tube current (mA), as listed in Table 1. Radiomic features extracted from tumors scanned with each option listed in Table 1 were compared against the reference values.

**Table 1:** Overview of the additional scan options included in this study. These options were chosen due to their common application in the clinic. *This is the kernel for the reference protocol.

| Additional Scanning Techniques |  |
| --- | --- |
| **Convolution Kernel** | *Standard, Lung & Bone Kernels |
| **Adaptive Statistical Iterative Reconstruction (ASiR) & ASiR-V** | 10, 20, and 30% ASiR and 10, 20, 30% ASiR-V |
| **Tube Potential (kVp)** | 100 kVP |
| **Tube Current (mA)** | 100 mA |

### 1.3 Radiomic feature extraction

The scanned tumors were contoured by an experienced radiologist using ImageJ software [26], as seen in Figure 2c. The contours were then propagated to all additional scans. Prior to radiomic feature extraction, the scans were re-sampled to an isotropic resolution of 1.0 mm. The computational environment for radiobiological research (CERR)[27] was used to extract 7 groups of radiomic features from each tumor: 22 first order (intensity) statistical measures[2, 28], 25 Grey-level co-occurrence matrix (GLCM)[2, 29], 16 gray-level run-length matrix (GLRLM), 5 neighborhood gray tone difference matrix (NGTDM)[28], 22 neighborhood gray level difference matrix (NGLDM), 16 Grey level size zone based features (GLSZM), and 2 peak and valley features were extracted from each tumor. These features are further described in the reference Zwanenburg, A, et al. [28]. The average value of each texture feature was computed over all 13 directions to obtain rotationally invariant features. For first order statistical features, a bin width of 25 was used. The GLCM features were extracted using a bin width of 10 and patch wise volume of 2 × 2 × 2.

For the second part of the study, the phantom was re-scanned, 3 times, without movement between scans, using the options listed in Table 1. The reference protocol scanning parameters were adjusted as needed. The reduced tube potential (100 kVp) and reduced tube current (100 mA) were acquired at reduced doses. Figure 5 shows scans of the radiomic phantom with each additional scan setting.

**Fig. 3.**
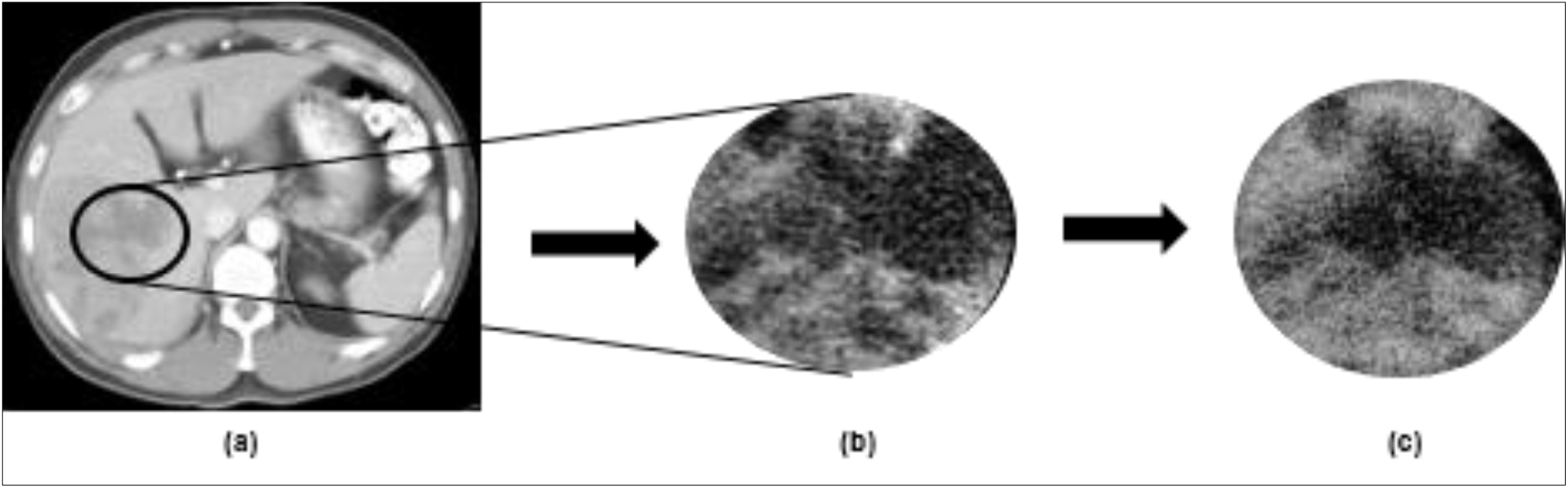
Comparison CT scan of segmented liver used as the background of the phantom with the resulting 3D print. (a) An axial slice from the patient CT showing the region of interest (ROI) around the heterogenous hepatic tissue. (b) The portion of the patient liver that the 3D print was modeled after. (c) A CT scan of the 3D print.

**Fig. 4.**
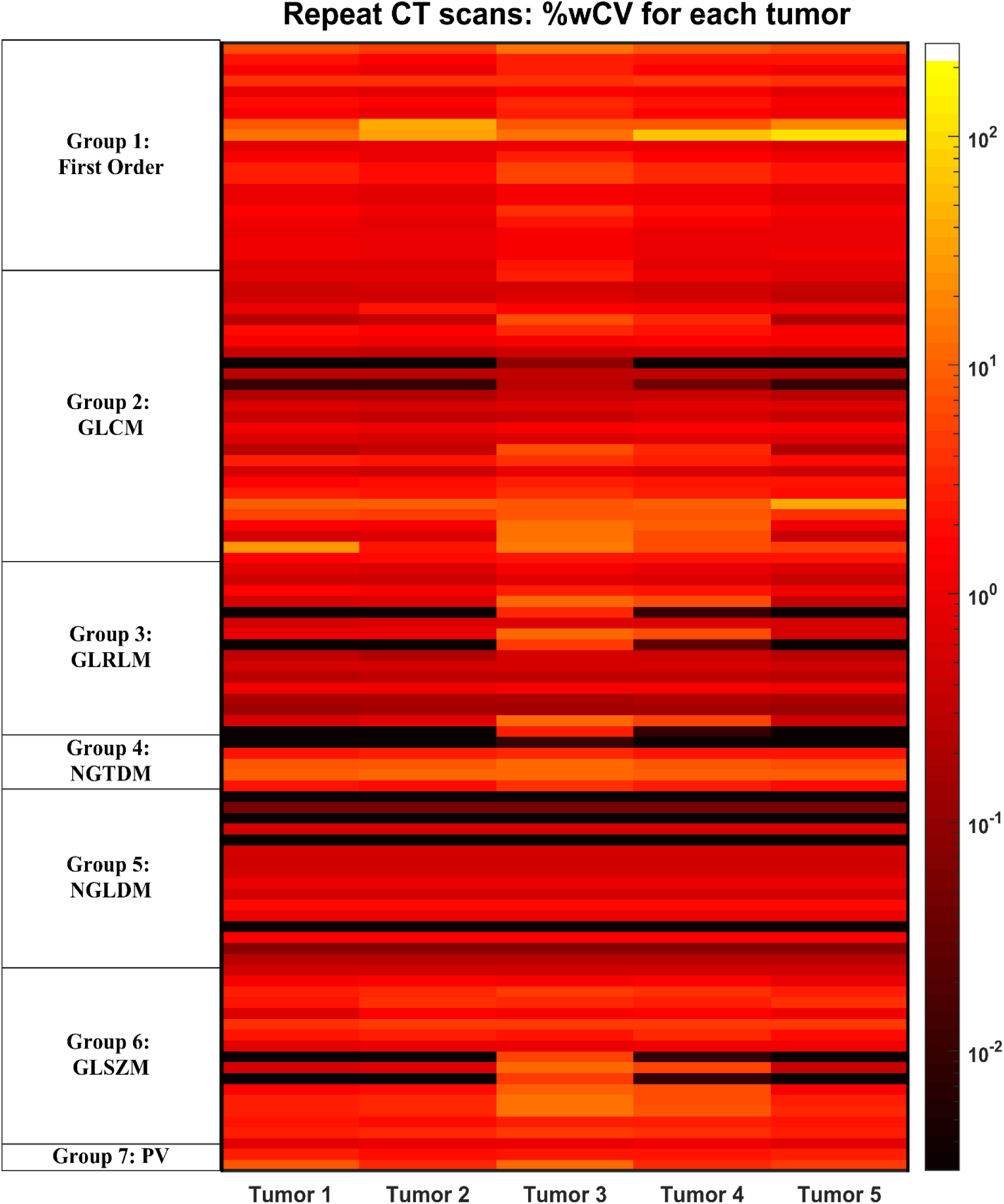
Within-subject coefficient of variation (%wCV) heat map of radiomic features for each tumor. The wCV was computed from the 32 repeat CT scans acquired with the reference protocol. The color map intensity is displayed on a logarithmic scale. The lighter the color, the lower the % wCV.

**Fig. 5.**
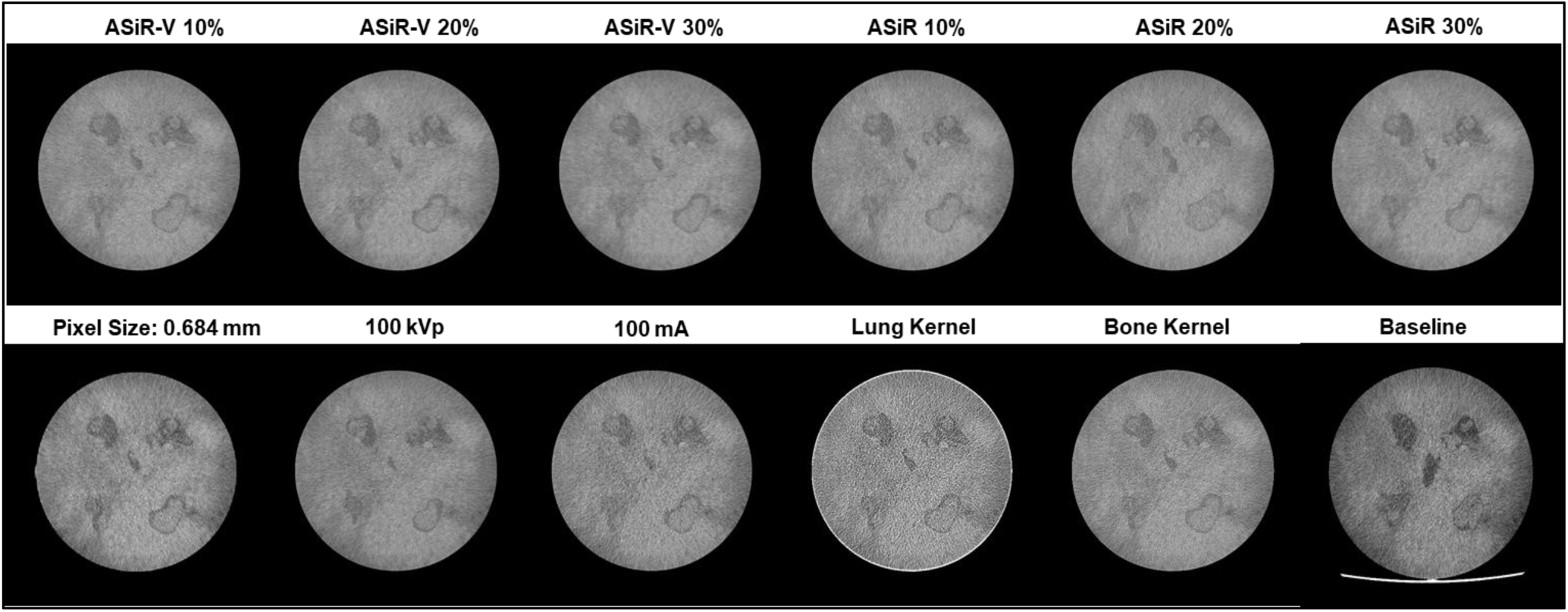
Cross-sectional CT images of the heterogenous and anatomical replicate tumors immersed within the radiomic phantom. It is displayed for each of the 11 different scan parameters that were used in this study. A window width of 40 and window level of 100 was used for all images. ASiR: Adaptive Statistical Iterative Reconstruction.

From CT scans of the phantom with each setting described above, a subset of radiomic features that have been found to be repeatable and prognostic for NSCLC (energy, grey level nonuniformity) and PDAC [2, 30] (entropy, energy, contrast and dissimilarity) were computed from each tumor. These features were then evaluated for their deviation from reference values, which were determined in the first part of this study.

### 1.4 Statistical Methods

Shown in Figure 3 is a slice of the cirrhotic liver (Figure 3a,b) that the background of the 3D print was modeled after. The structural similarity index (SSIM) [31] was used to calculate the similarity between the original patient image and the resulting 3D print. Prior to the comparison, the patient DICOM images were converted into png’s. A detailed derivation and explanation of SSIM can be found by Wang, Z, et al. [31].

Repeatability (i.e. precision) of radiomic features was evaluated [7] using the within-subject coefficient of variation (wCV, %).

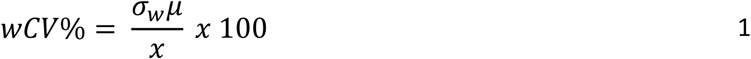

where *σ*_*w*_ is the within-subject standard deviation and *μ* is the mean of individual radiomic features. A *wCV*% less than 10% was considered as being repeatable. The 95% confidence interval (CI) for the *wCV* was calculated using chi squared (*x*^2^) as the pivotal statistic as follows:

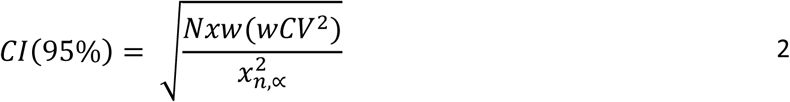

where N is the number of tumors, 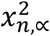 is the percentile of the distribution with n degrees of freedom. The lower bound, α is 0.975 and the upper bound α is 0.025.

To determine the deviation of select radiomic features from reference values, the percent deviation (pDV, %) was calculated as follows:

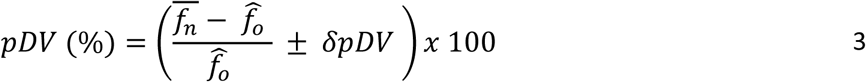

where 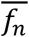 is the average value of the radiomic feature extracted from images of each tumor across the different scanning parameters, and 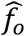 is the average of the reference value as described above.

## Results

### 1.6. Repeatability

The SSIM between the real patient image (Figure 3b) and the 3D print (Figure 3c) was found to be SSIM = 0.71. A SSIM value closer to 1 suggests more similarity. Figure 4 is a heat map showing the wCV’s (%) for the 104 radiomic features computed from the 30 repeat scans of the phantom. Within the heatmap, the darker the color, the lower the wCV. Out of the 104 radiomic features, 87 were found to have a wCV < 10%. Group 5 radiomic features, the neighboring gray level dependence matrix (NGLDM), which are characterized as being rotationally invariant, and those that capture the coarseness of overall texture within an image were the only feature class with wCV < 10% across all tumors included in the phantom.

Table 2 summarizes those radiomic features with wCV (%) values that exceeded 10%.. As shown in Table 2, the repeatability of some features with wCV (%) > 10% depended on tumor type. For example, when extracted from images of the 2^nd^ and 5^th^ tumor, FO8: skewness had a wCV = 33.09 % (CI: 26.01 to 42.42%) and wCV = 23.57% (CI: 18.70 to 26.01%), but wCV < 10% when it was calculated from tumors 1 (NSCLC), 3, and 4 (PDAC). GLCM 25: first information correlation also had a wCV > 20% only for the 1^st^ tumor. Two of the 17 radiomic features, FO9: kurtosis and NG4: complexity were found to have wCV > 10% for all tumors. The non-repeatable features were excluded from any further analysis.

**Table 2:** Radiomic features with the within-subject coefficient of variation (% wCV) > 10% and groups with wCV (%) < 10%.

| wCV < 10 % |  | wCV > 10% |  |
| --- | --- | --- | --- |
| Group 1 | 14 out of 22:<br>FO 2 to 7, & 10 to 22 | FO1: Min | T1 |
|  |  | FO8: Skewness | T2, T5 |
|  |  | FO9: Kurtosis | T1-T5 |
| Group 2 | 22 out of 26 features:<br>GLCM 1 to 20, 22, & 26 | GLCM21: Cluster Shade | T1, T2, T4, T5 |
|  |  | GLCM23: Haralick Correlation | T3, T4 |
|  |  | GLCM24: Auto Correlation | T3 |
|  |  | GLCM25: First Information Correlation | T1, T3 |
| Group 3 | 13 out of 16 radiomic<br>features:GLRLM1: 1 to 6;<br>GLRLM 8 to 14, & 16. | GLRLM14: High gray level run emphasis | T3 |
|  |  | GLRLM17: Long run high gray level<br>emphasis | T3 |
|  |  | GLRLM15: Short run high gray level<br>emphasis | T3 |
| Group 4 | 3 out of 5:<br>NGTDM: 1, 2 & 5 | NGTDM3: Busyness | T3 |
|  |  | NGTDM4: Complexity | T3 |
| Group 6 | 12 out of 16:<br>GLSZM: 1 to 8, 10, & 14 to<br>16 | GLSZM9: High gray Level zone emphasis | T3 |
|  |  | GLSZM11: Small area high gray level<br>emphasis | T3 |
|  |  | GLSZM12: Large area low gray level<br>emphasis | T3 |
|  |  | GLSZM13: Large area high | T3 |
| Group 7 | 1 out of 2: Peak | PV2: Valley | T3 |

### 1.7 Deviation from Baseline QIB values

For subjective comparisons, Figure 5 shows representative cross-sectional slices of the phantom for each parameter in Table 1. The pDVs (%) for energy and grey level nonuniformity (GLNU), which are prognostic for NSCLC, are shown in Figure 6. Both features had pDV (%) values that were negative. For ease of viewing, the plot was inverted. Except for a larger pixel size (PS, i.e. DFOV) of 0.684 mm, the pDV (%) was < -30% across all other scan parameters. The negative pDV (%) implies that the radiomic features were less than the reference values.

**Fig. 6.**
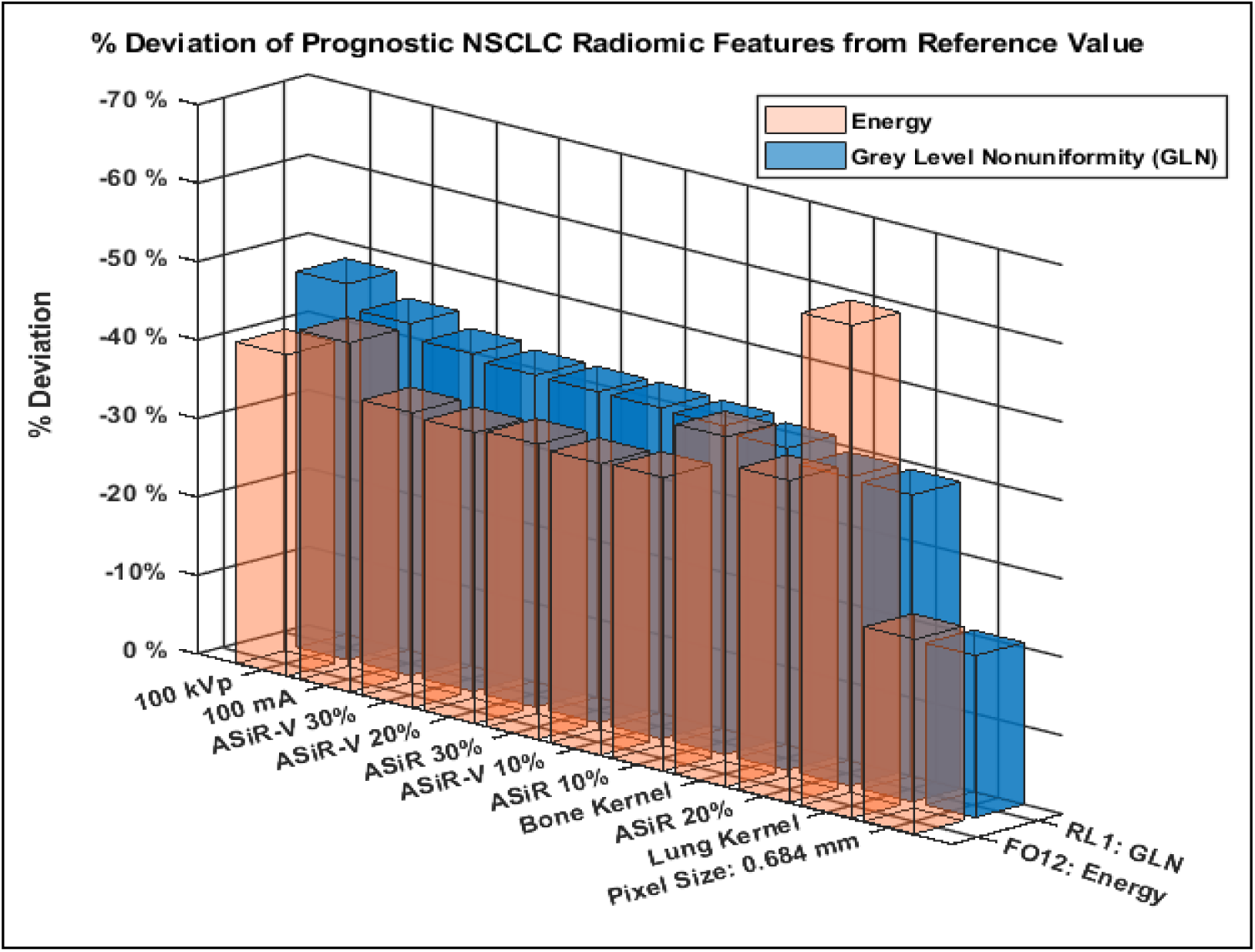
The percent deviation (pDV, %) of prognostic radiomic features for Non-Small Cell Lung Carcinoma (NSCLC). Left: Energy, and Right: Grey level nonuniformity texture features. The darker colors indicate lower pDV (%).

The boxplots in Figure 7 show the total deviation of prognostic PDAC radiomic features across all scan settings. Except for the 5^th^ PDAC tumor, the pDV (%) distribution for energy remained within pDV (%) ± 20% across most scan parameters. The interquartile range for entropy, contrast, and dissimilarity were found to depend on tumor type.

**Fig. 7.**
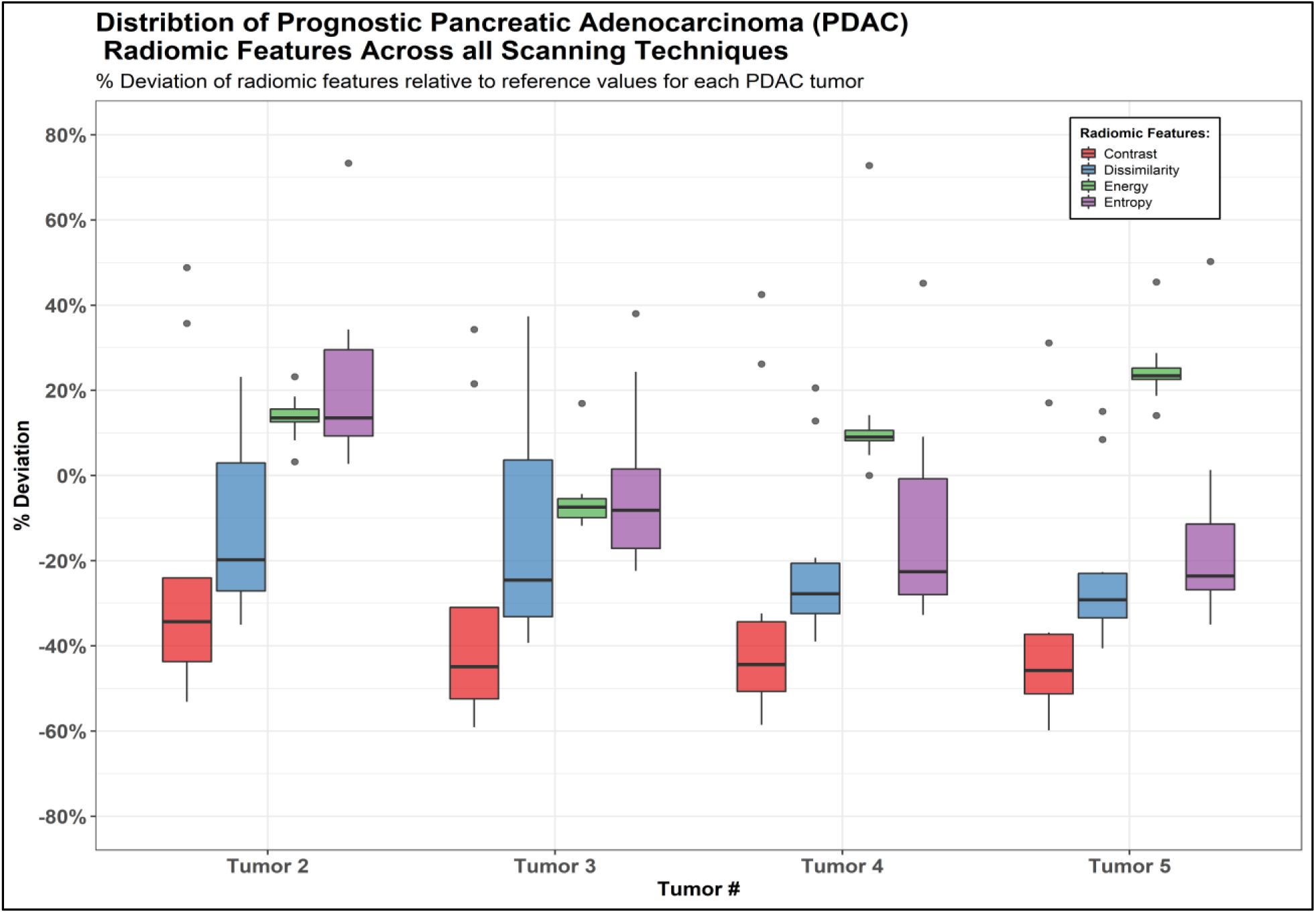
Box plots showing the deviation of prognostic PDAC radiomic feature values: contrast, dissimilarity, energy and entropy (compared with the reference values) as a function of tumors imaged across all scanning techniques. Displayed is the median, interquartile range (25^th^, 75^th^), maximum, minimum and outliers.

Figure 8 show a more granular assessment of radiomic feature deviation from the reference value as a function of scanning technique. The pDV(%) for each radiomic feature value was found to depend on tumor type and the imaging condition. Contrast, which assesses the variations in grey levels within an ROI [28], had pDV’s (%) that exceeded ± 20 % for most scanning parameters across each of the tumors. When a reduced tube current of 100 mA was used, the pDV (%) for contrast was < 20% when measured from the 5^th^ tumor. In addition, across all tumors contrast was calculated to have a pDV (%) of 100% with application of the lung kernel. However, only for the 2^nd^ and 3^rd^ tumor, contrast had a pDV(%) = 100% when ASiR 20% was used to reconstruct images. Dissimilarity is similar to contrast in that it sums the difference of discretized grey level intensity values from the GLCM, whereas contrast sums the squared difference of discretized intensity values[28]. Except for the lung kernel where pDV (%) = 100%, the trends in pDV (%) for other scanning parameters were similar to contrast, but less pronounced.

**Fig. 8.**
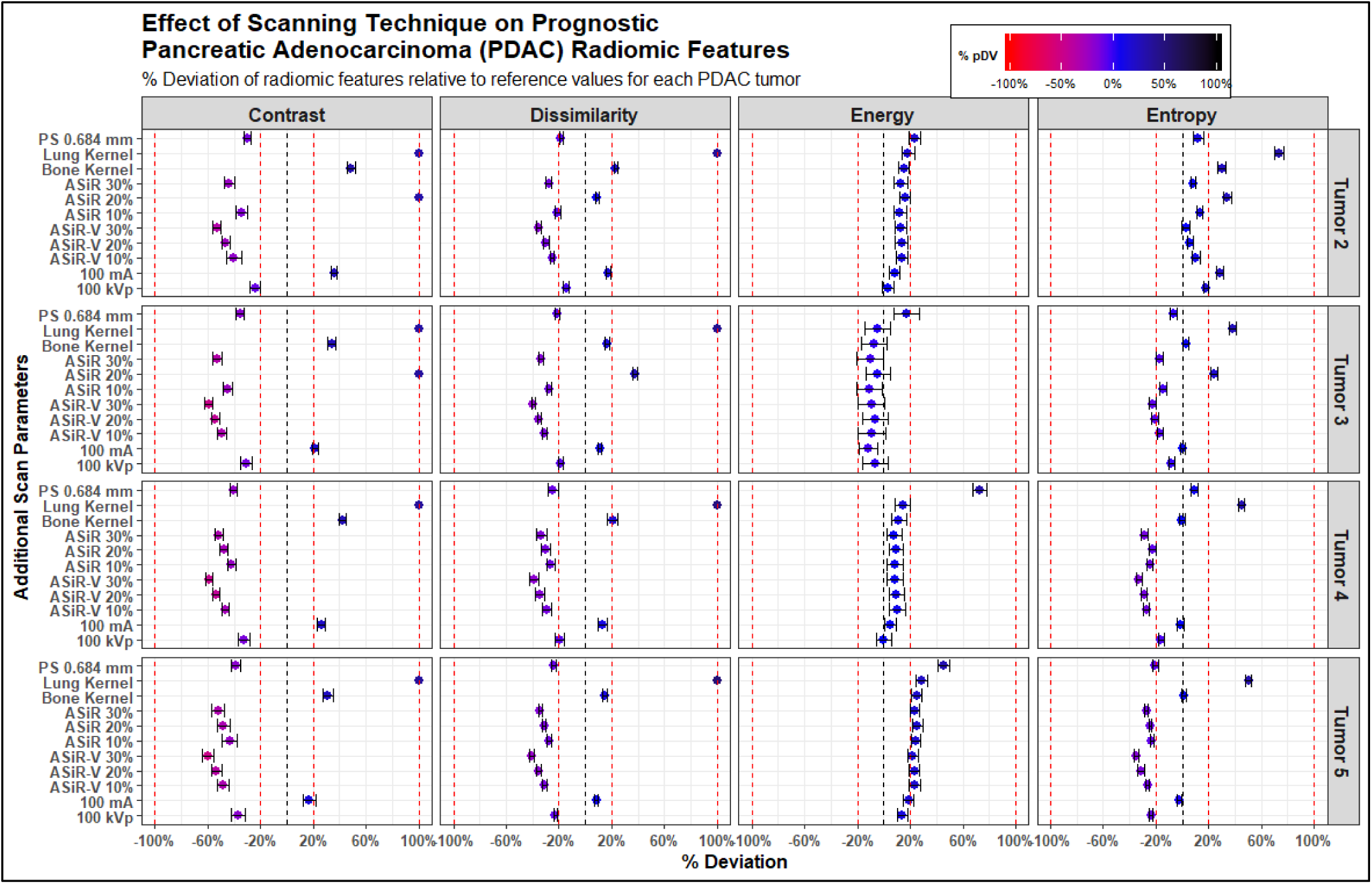
Dot plots that show the percent deviation (pDV, %) of quantitative radiomic feature values as a function of each scanning technique and tumor type. Lower, red values indicate a negative pDV (%). Darker points indicate larger positive pDV (%). The blue points are pDV (%) values close to zero. The vertical dotted lines indicate the ± 0%, 20% and 100% pDV’s (%).

With a reduced tube current of 100 mA and tube potential of 100 kVp, the pDV (%) for energy, which is a measure of the overall tumor volume density, was slightly within ± 20% when derived from the 5^th^ tumor. Only for the 4^th^ tumor, the pDV (%) for energy was > 60% when the PS = 0.684 mm. For the remaining tumors, the energy remained within ± 20% across sanning parameters. Across tumors 3, 4, and 5, the pDV (%) for entropy remained within ± 3% when the a reduced tube current and the bone kernel were used.

## Discussion

Here we describe a process to manufacture anatomically realistic 3-D printed tumors embedded in heterogenous backgrounds for identifying and evaluating robust quantitative radiomic features. Capturing the anatomical structure and reproducing perceptually similar attenuation patterns of tumors seen on CT scans will allow us to extract radiomic features that can serve as “ground truth” and as a result, piece together and reduce sources of variability between scanners, and acquisition protocols/parameters. Furthermore, computing the acceptable range of variation from patient studies consists of unquantifiable errors or sources of uncertainty. From the kinetic behavior of contrast media to the limitation of using only one scanner at a single institution[7, 10, 28], using patients for determining the acceptable range of variation is not practical. Similarly, homogenous phantoms that are vastly different from abnormal and normal human tissue seen on CT scans, will not be able to assess how the background of a lesion can affect radiomic features calculated from tumors [7, 10, 28]. This 3D printed phantom affords the opportunity to filter out quantitative radiomic features that are sensitive to CT scanning parameters and potentially allows one to establish reference ranges for the robust quantitative radiomic features [21].

An important finding from this experiment is that radiomic features were dependent on the location and/or type of tumor. This finding is essential to take into consideration when using quantitative radiomic features to characterize metastases lesions. A feature that can robustly describe the primary tumor site may not hold the same meaning for distant metastatic disease. This finding is conceptually similar to the understanding that metastasized lesions may not respond to treatment used for the primary tumor site [32].

Prior studies that used homogenous nodules or homogenously textured phantoms that found radiomic features to be repeatable might under or overestimate findings. In addition, several research efforts have generated prognostic radiomic signatures for a variety of disease types [9, 17]. Without established reference ranges for invariant quantitative radiomic features that establish thresholds below or above which disease is present, false correlations may ensue. For example, from Group 3, GLRLM feature long run high gray level emphasis (LRHGLE) was found to be a good discriminator of renal cell carcinomas (RCC) from other RCC subtypes [33]. We find that the repeatability of LRHGLE depends on tumor type, where the wCV exceeded 10% for the 3^rd^ tumor (wCV (%) = 14.27% (95% confidence interval (CI): 11.40% to 19.10%), but was < 1% for tumors 1, 2 and 5, and 7.30% (CI: 5.86% to 9.83%) for the 4^th^ tumor. In another study[9, 16], first order (FO9) kurtosis, from the group 1 radiomic feature, was found to be a prognostic measure of disease free survival after the end of radiation therapy for NSCLC [9, 16]. Based on our repeatability results, kurtosis presented with wCV (%) > 10% for all tumors in the phantom. However, a wCV (%) exceeding 10% was arbitrarily chosen to indicate a variable quantitative radiomic feature. Within this context, anatomically realistic 3D printed phantoms are a promising alternative to patients when trying to determine acceptable ranges of variation and the uniform reporting of repeatability and reproducibility of quantitative radiomic features.

Sensitivity to varying CT scan parameters can lead to false positives or negatives and must be quantified prior to the clinical implementation of radiomic metrics [17, 19, 23, 34]. The drive towards automation has resulted in CT scanning parameters that automatically adjust for a patient’s body habitus and attenuation characteristics. Using clinically relevant scanning parameters to acquired additional scans of the phantom, we again find that the deviation of previously discovered prognostic radiomic features depend on tumor type and the applied scanning method.

There are several limitations to this work. The material used for 3D printing is limited in its HU value range. It cannot recreate all HU values seen in patient CT exams. As such, the change in intensity values with changes in the X-ray beam spectrum resulting from use of different bowtie filters, patient off centered positioning, different tube potential’s, etc. may exacerbate the amount of deviation or variation of a radiomic feature. However, with this 3D printed phantom, the end goal is to eliminate those radiomic features that would be sensitive to such changes, even if the lack of repeatability or reproducibility would be overstated. Even though the lesions and background of the phantom were modeled after real tumors seen on CT scans, the lack of surrounding tissue and bony structure typically found within the abdomen reduces the degree to which a one to one correlation can be made. Future works will aim to include the fat content and bony structure seen on patient CT scans. Since the phantom was a static object, we did not incorporate motion into this study, which is an essential element to consider. Published methods that use motorized devices could be considered in future studies. Since the purpose of this study was to demonstrate the feasibility and potential role of a patient informed 3D printed phantom, statistical significance was not evaluated in this study. In addition, the spectral characteristics of CT scanners, calibration, electronic noise unique to CT scanner were not assessed. These sources of uncertainty could be the culprits that bias results and without accounting for their influence on quantitative radiomic metrics, results might be falsely discrepant from ground truth values.

A final critical element of this phantom is its unique ability to determine the influence of manual, semi-automated and automated segmentation methods on radiomic feature stability. Due to the amount of data collected, evaluation of segmentation methods was left to future works.

## Conclusion

A first step in any quantitative radiomic feature discovery pipeline will need to be robustness studies that are conducted with anatomically realistic phantom. The reproduction of diseased tissue morphology and contrast differences in a patient informed phantom with background cirrhotic liver was demonstrated using the novel voxel-based 3D printing method. These realistic phantoms can be used to delineate invariant radiomic features. These robust features can then be investigated for their potential clinical application.

## Acknowledgements

This research was funded in part through National Institutes of Health/National Cancer Institute Cancer Center Support grant P30 CA008748. The authors thank Brent Chanin from MediPrint, LLC for his consultative help in designing the lung model.

## Footnotes

a Dots Per Inch: DPI

## References

1. Choi, E.-R., et al. Quantitative image variables reflect the intratumoral pathologic heterogeneity of lung adenocarcinoma. 2016. 7(i41): p. 67302.

2. Aerts, H.J., et al. Decoding tumour phenotype by noninvasive imaging using a quantitative radiomics approach. 2014. 5: p. 4006.

3. Bak, S.H., et al. Prognostic Impact of Longitudinal Monitoring of Radiomic Features in Patients with Advanced Non-Small Cell Lung Cancer. 2019. 9(1): p. 8730.

4. Raman, S.P., et al. CT texture analysis of renal masses: pilot study using random forest classification for prediction of pathology. 2014. 21(12): p. 1587–1596.

5. Grove, O., et al. Quantitative computed tomographic descriptors associate tumor shape complexity and intratumor heterogeneity with prognosis in lung adenocarcinoma. 2015. 10(3): p. e0118261.

6. Aerts, H.J., et al. Defining a radiomic response phenotype: a pilot study using targeted therapy in NSCLC. 2016. 6: p. 33860.

7. Raunig, D.L., et al., Quantitative imaging biomarkers: a review of statistical methods for technical performance assessment. 2015. 24(1): p. 27–67.

8. Zhao, B., et al., Reproducibility of radiomics for deciphering tumor phenotype with imaging. 2016. 6: p. 23428.

9. Avanzo, M., J. Stancanello, and I.J.P.M. El Naqa, Beyond imaging: The promise of radiomics. 2017. 38: p. 122–139.

10. Sullivan, D.C., et al., Metrology standards for quantitative imaging biomarkers. 2015. 277(3): p. 813–825.

11. Solomon, J., et al., Comparison of low-contrast detectability between two CT reconstruction algorithms using voxel-based 3D printed textured phantoms. 2016. 43(12): p. 6497–6506.

12. Solomon, J. and E.J.M.p. Samei, Quantum noise properties of CT images with anatomical textured backgrounds across reconstruction algorithms: FBP and SAFIRE. 2014. 41(9): p. 091908.

13. Solomon, J., J. Wilson, and E.J.M.p. Samei, Characteristic image quality of a third generation dual-source MDCT scanner: Noise, resolution, and detectability. 2015. 42(8): p. 4941–4953.

14. Obuchowski, N.A., et al., Statistical issues in the comparison of quantitative imaging biomarker algorithms using pulmonary nodule volume as an example. 2015. 24(1): p. 107–140.

15. Kessler, L.G., et al., The emerging science of quantitative imaging biomarkers terminology and definitions for scientific studies and regulatory submissions. 2015. 24(1): p. 9–26.

16. Huang, Y., et al., Radiomics signature: a potential biomarker for the prediction of disease-free survival in early-stage (I or II) non—small cell lung cancer. 2016. 281(3): p. 947–957.

17. Lambin, P., et al., Radiomics: the bridge between medical imaging and personalized medicine. 2017. 14(12): p. 749.

18. Shafiq-ul-Hassan, M., et al., Intrinsic dependencies of CT radiomic features on voxel size and number of gray levels. 2017. 44(3): p. 1050–1062.

19. Mackin, D., et al., Harmonizing the pixel size in retrospective computed tomography radiomics studies. 2017. 12(9): p. e0178524.

20. Bader, C., et al., Making data matter: Voxel printing for the digital fabrication of data across scales and domains. 2018. 4(5): p. eaas8652.

21. Samei, E., et al., Design and fabrication of heterogeneous lung nodule phantoms for assessing the accuracy and variability of measured texture radiomics features in CT. 2019. 6(2): p. 021606.

22. Zhao, B., L.H. Schwartz, and M.G.J.T.C.I.A. Kris, Data from RIDER_Lung CT. 2015.

23. Zhao, B., et al., Evaluating variability in tumor measurements from same-day repeat CT scans of patients with non–small cell lung cancer. 2009. 252(1): p. 263–272.

24. Clark, K., et al., The Cancer Imaging Archive (TCIA): maintaining and operating a public information repository. 2013. 26(6): p. 1045–1057.

25. Newell Jr, J.D., J. Sieren, and E.A.J.J.o.t.i. Hoffman, Development of quantitative CT lung protocols. 2013. 28(5).

26. Schneider, C.A., W.S. Rasband, and K.W.J.N.m. Eliceiri, NIH Image to ImageJ: 25 years of image analysis. 2012. 9(7): p. 671.

27. Deasy, J.O., A.I. Blanco, and V.H.J.M.p. Clark, CERR: a computational environment for radiotherapy research. 2003. 30(5): p. 979–985.

28. Zwanenburg, A., et al., Image biomarker standardisation initiative. 2016.

29. Haralick, R., K. Shanmugam, and I.J.I.T.S.M.C. Dinstein, Textural features for image analysis. 1973. 6: p. 610–621.

30. Cozzi, L., et al., Computed tomography based radiomic signature as predictive of survival and local control after stereotactic body radiation therapy in pancreatic carcinoma. 2019. 14(1): p. e0210758.

31. Wang, Z., et al., Image quality assessment: from error visibility to structural similarity. 2004. 13(4): p. 600–612.

32. Yoo, B., B.C. Fuchs, and Z.J.F.i.o. Medarova, New directions in the study and treatment of metastatic cancer. 2018. 8.

33. Yu, H., et al., Texture analysis as a radiomic marker for differentiating renal tumors. 2017. 42(10): p. 2470–2478.

34. Boedeker, K.L., et al., Emphysema: effect of reconstruction algorithm on CT imaging measures. 2004. 232(1): p. 295–301.

